# MiR-590-5p sensitises pancreatic ductal adenocarcinoma cells by blocking autophagy via targeting ATG3

**DOI:** 10.1101/548610

**Authors:** Fazhao Li, Jun He, Susun Liu, Yawei Zhang, Leping Yang

## Abstract

Radio-resistance is a growing concern in treating patients with pancreatic cancer (PC). Here we investigated the role of miR-590-5p in the radio-resistance of PC cells. We developed radioresistant PC cell lines and followed by microarray analysis and levels of miRs compared to parental cell lines. PC cells were transfected using either miR mimics or inhibitors followed by clonogenic survival assays. We also studied the effect of miR-590-5p on autophagy using electron microscopy and immunoblot analysis. In addition, the luciferase assay was used to identify potential targets. The radio-resistant PC cells exhibited decreased expression of miR-590-5p, with elevated autophagy against the parental cells. The over-expression of miR-590-5p inhibited radiation-mediated autophagy, while inhibitors induced autophagy in PC cells. The up-regulation of miR-590-5p enhanced the radio-sensitivity of PC cells. We confirmed ATG-3 as a target of miR-590-5p, whose levels were unregulated in radio-resistant cells. We also found that levels of ATG-3 were associated with autophagy. Expression of miR-590-5p inhibited radiation-mediated autophagy and enhanced the radio-sensitivity of PC cells.

## 1. Introduction

Pancreatic cancer (PC) has been confirmed to be one of the leading causes of mortality, with an overall survival rate of less than 5% [1]. Surgery, which has a poor success rate, is currently the only major treatment [2]. It has been reported that a large population of PC cells in locally advanced pancreatic cancer are involved in radioresistance when PC patients are treated with radiation therapy [1]. However, due to the inherent chemoresistance and radioresistance of PC, the prognosis is poor, and surgery usually fails to produce a positive outcome [3,4]. Radioresistance is therefore a major element of PC treatment, and understanding the mechanism of radioresistance process at a molecular level may lead to treatment improvements, as well as better therapeutic outcomes.

To date, studies have identified some links between radioresistance and genes specifically involved in PC, which play a role in DNA damage and enhancement of DNA repair [5–7]. These studies have helped elucidate the mechanisms involved in radiosensitivity, but the specific details of the pathways and molecular mechanisms involved in resistance remain unknown. MicroRNAs (miRs) are small non-coding RNA molecules of 22 nucleotides, which are responsible for regulating the expression of genes at the posttranscriptional level. The miRs bind either completely or partially to mRNA, altering the expression of proteins either by inhibiting translation or by degradation of mRNA [8]. Numerous studies have suggested a role of miRs in many human disorders, including cancer [9]. Some miRs are tumour suppressors, while others modulate the expression of tumour suppressor genes [10]. A number of miRs have been confirmed to be associated with improved survival of patients, and may be useful when designing effective treatments involving radiation and chemotherapy [11–13]. However, limited data are available regarding the expression of miRs, and the mechanisms and functions of miRs remain relatively unexplored.

In the present investigation, we showed that miR-590-5p enhanced the radiosensitivity of PC cells by inhibiting radiation-mediated autophagy. We also confirmed that miR-590-5p targeted autophagy-related protein-3 (ATG-3), which was upregulated in PC cells and positively correlated with autophagy and radiation sensitivity. Overall, we identified the potential role of miR-590-5p in autophagy and confirmed that miR-590-5p increased the sensitivity of PC cells to radiation by inhibiting radiation-mediated autophagy.

## 2. Materials and Methods

### 2.1. Subjects and tissue samples

Human PC tissue specimens were obtained from 54 patients at the Pancreatic and Gallbladder Surgery Ward, The Second Xiangya Hospital of Central South University, Changsha, Hunan, China by surgical removal. The study objectives and protocol were explained to all patients, who signed informed consent forms, and the study was approved by the institutional research committee at the same institution. The patient characteristics were recorded and are listed in Table 1. All patients were treated with surgical resection, but were not subjected to any type of chemotherapy or radiotherapy. The tumours and lymph nodes obtained after surgical resection were histologically examined using haematoxylin and eosin staining and the TNM classification. Histological examination showed that all tumours were invasive ductal adenocarcinomas. The resected cancerous tissues were frozen, fixed with formalin, and embedded in paraffin to obtain tissue sections, and the sections were immunochemically analysed for the expression of miR-590-5p.

**Table 1:**
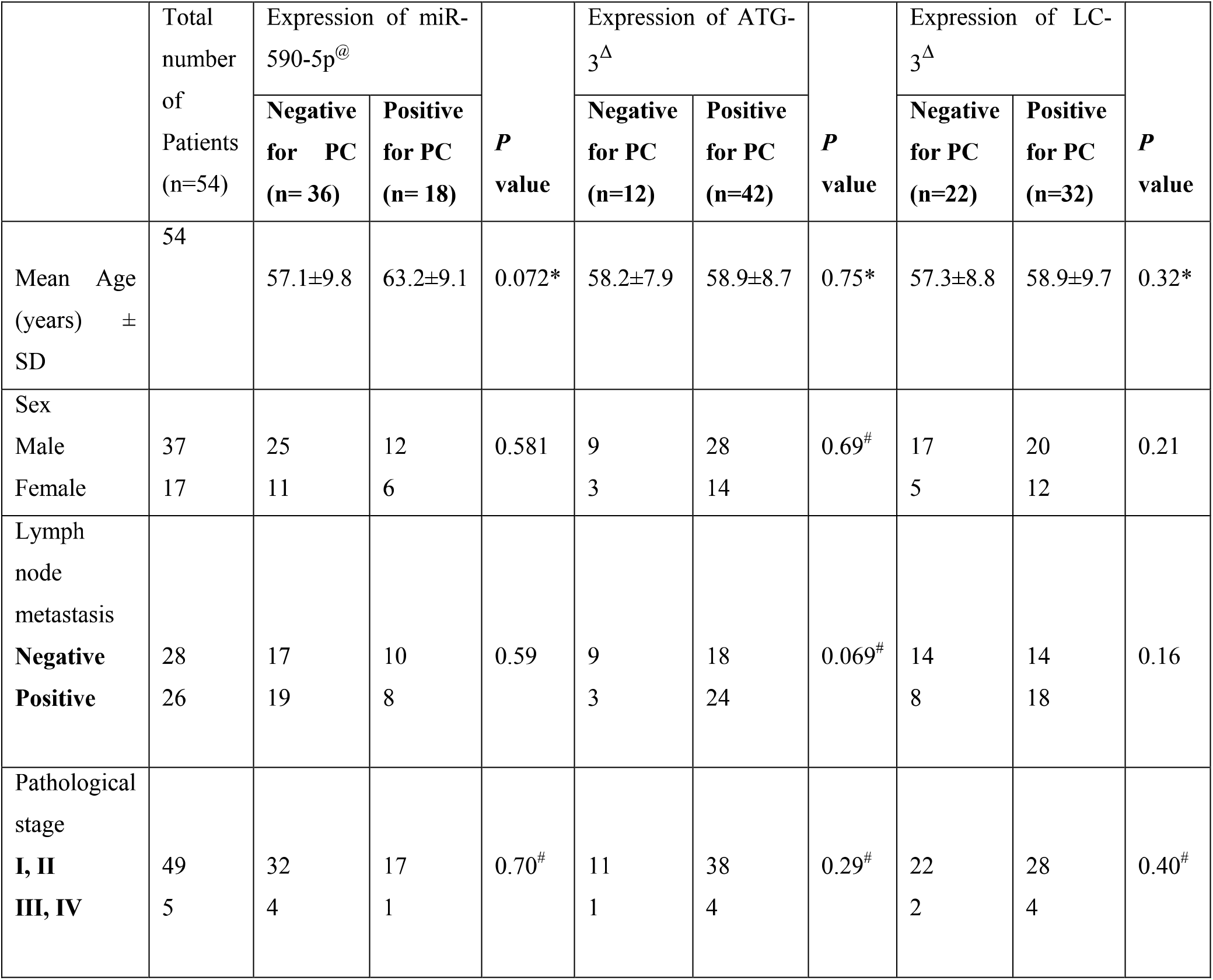
Patient data, indicating correlation between pathological features of pancreatic cancer patients and levels of LC-3, miR-590-5p and ATG-3. @ = levels of miR-590-5p, detected by in situ hybridization, Δ = levels of ATG-3 and LC-3 detected by immunohistochemical analysis. ^**b**^Compared and established using student *t* test ^#^Compared with the help of Fisher exact test. Rest of the *P* values was calculated using Pearson x^2^ test.

### 2.2. Human PC cells and animals

The AsPC-1 and Capan-2 human PC cell lines were obtained from Sigma-Aldrich (St. Louis, MO, USA) and Thermo Fisher (Waltham, MA, USA), respectively, and were cultured in growth medium as directed by the suppliers. Both cell lines were maintained under humid conditions at 37°C and 5% CO_2_. The cell lines were tested regularly for the presence of any mycoplasma contamination. For *in vivo* studies, female BALB/c nude mice (5–6 weeks of age) were obtained from the Animal Department of The Second Xiangya Hospital of Central South University, Changsha, Hunan, China. The experimental protocol was approved by the institutional animal ethics committee of the same institution, and all experiments adhered to guidelines of the “Animal Protection Law of the People’s Republic of China-2009” (approval number: TSXH/AEC/2018/PC11). All animals were housed under controlled conditions with laminar air flow, and were provided with water and food ad libitum.

### 2.3. Establishment of radioresistant cancer cell sublines

The radioresistant cancer cell sublines, AsPC-1/RR1, AsPC-1/RR2, and Capan-2/RR2, were produced from parental cell lines AsPC-1 and Capan-2. The AsPC-1 and Capan-2 cell lines were seeded in T25 flasks at a density of 1 × 10^6^ cells in culture medium. Upon reaching 70% confluence, the cells were exposed to ^60^Co radiation at 2 Gy/min, and upon reaching 80% confluence, the cells were treated with trypsinisation reagent (Thermo Fisher) and were cultured again in a new flask. Upon reaching 70% confluence, the cells were subjected to irradiation of increasing concentrations (4, 6, 8, and 10 Gy/min). For the study, two clones/cell line were chosen and labelled as AsPC/RR1, AsPC/RR2, Capan-2/RR1, and Capan-2/RR2. These clones were again exposed to five cycles of 10 Gy/min radiation, and the cells showing radioresistance were cultured in medium and used as radioresistant human PC cell lines in subsequent experiments.

### 2.4. Colony formation assay

The PC cells were cultured at a density of 1 × 10^6^ cells in Gibco cell culture medium (Thermo Fisher) in flasks. After 24 h, the cells were exposed to radiation (0, 2, 4, 6, 8, or 10 Gy/min) once and seeded into 6-well plates, followed by a 2-week incubation without replacing the medium. The cell cultures were then fixed in methanol and stained with crystal violet. Using a light microscope, the number of colonies containing > 50 cells were counted. Survival fraction (SF) was used as an index of colony formation, and was calculated using the equation: SF = number of colonies/number of cells seeded × (plating efficiency/100). The survival curve was then constructed using a multi-target single hit model.

### 2.5. Isolation of RNA, microarray analysis, and semi-quantitative reverse transcription-polymerase chain reaction (RT-PCR)

The RNA of radioresistant cells was isolated using the TRIzol reagent (Thermo Fisher) following the supplier’s instructions. The RNAs were extracted using the RNeasy Mini Kit (Qiagen, San Diego, CA, USA). The arrays were performed using the TargetScan prediction algorithm (http://www.targetscan.org/) for identifying targets of miR-590-5p. The TaqMan miRNA assay (Applied Biosystems, Foster City, CA, USA) was used according the manufacturer’s instructions, to measure the expression of mature mRNAs. To perform real-time PCR studies, miR-590-5p specific TaqMan primers were used to find the average cycle threshold value of each sample. Semi-quantitative real-time RT-PCR was performed to determine the relative expression of specific mRNAs. The primer sequences are listed in Table 2.

**Table 2:**
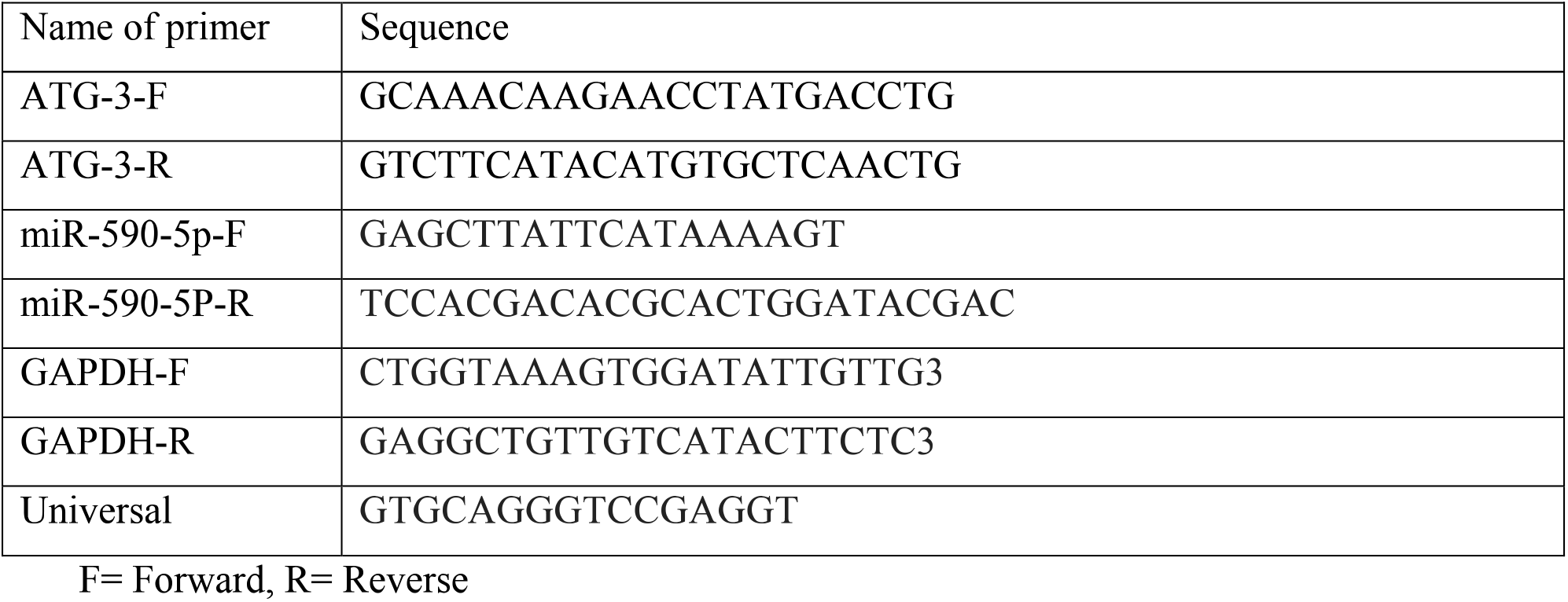
Sequence of primers used in the study

### 2.6. Immunoblotting studies

Immunoblotting was conducted to evaluate protein expression. The cultured cells were treated to extract proteins using a protein extraction kit (Thermo Fisher), with bovine serum albumin as the standard. The cytoplasmic proteins were also extracted from cells using the Nuclear & Cytoplasmic Extraction Kit (G-Biosciences, St. Louis, MO, USA). The proteins were resolved using sodium dodecyl sulphate-polyacrylamide gel electrophoresis (10%) and then transferred to nitrocellulose membranes (Thermo Fisher). The membranes were first blocked using 5% non-fat milk, then incubated with primary antibodies. Proteins bands were visualised using an Immun-star AP chemiluminescence kit (Bio-Rad, Hercules, CA, USA). The primary antibodies used for the experiment are listed in Table 3. The transfecting miR-590-5p mimics and inhibitors were obtained from Sigma-Aldrich, and the negative control (NC) oligonucleotides were nonhomologous to sequences of human genes. Lipofectamine 2000 reagent (Invitrogen, Carlsbad, CA, USA) was used in the transfection process.

**Table 3:**
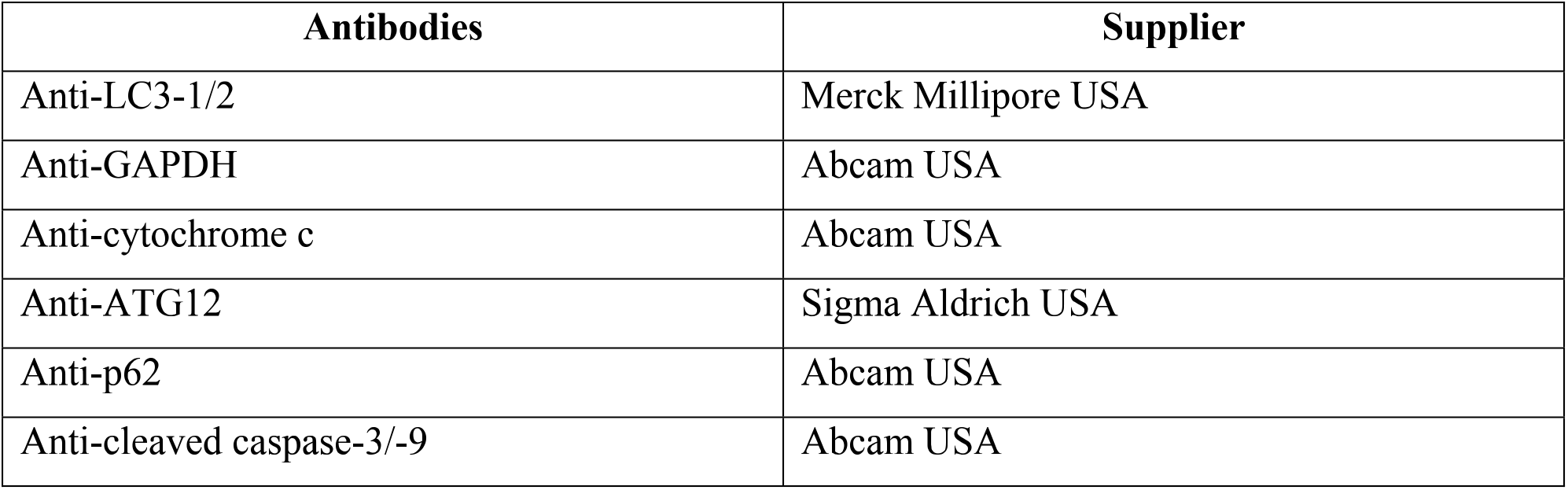
The primary antibodies used for the study

### 2.7. Vector construction

The lentiviral expression vector was subcloned using the partial pri-miR-590-5p gene. The amplification of ATG-3 was performed by cloning the sequence into the pCDHCMV-MCS-EF1-Puro lentiviral vector to obtain pCDH-ATG-3. Short RNAs (shRNAs) were produced using the pLKO-1-puro lentiviral vector. The wild-type (WT) 3’-untranslated region (UTR) segment of ATG-3 mRNA was amplified using PCR and inserted into the Renilla luciferase gene. Site-directed mutagenesis was performed according to the manufacturer’s instructions to generate 3’-UTR segments of WT ATG-3 mRNA. The primers used for the experiment are listed in Table 2.

### 2.8. Luciferase assay

AsPC-1 and Capan-2 cells were co-transfected using 80 ng of the pRL-TK-Renilla-luciferase reporter plasmid, 40 ng of plasmid, and 20 nmol/L of the indicated RNAs. A dual luciferase (firefly and Renilla) assay system (BPS Bioscience, San Diego, CA, USA) was used to quantify the luciferase activity 24 h after 3× transfections.

### 2.9. Electron microscopy

After treatment, the cells were fixed in 2.5% glutaraldehyde (Sigma-Aldrich) solution in 0.1 mol/L sodium cacodylate (Sigma-Aldrich). The tissue samples were fixed using 1% osmium tetraoxide and dehydrated using ethanol and propylene oxide; then, the tissues were sliced into 50 nm sections followed by staining with 3% uranyl acetate and lead citrate. An SY9000 electron microscope (Hitachi, Tokyo, Japan) was used to capture images.

### 2.10. Generation of stably expressing cell lines

The selected AsPC-1 and Capan-2 PC cell lines were transfected with the GFP-LC3 lentiviral vector (Invitrogen). Transfection flow cytometry analysis was then conducted to isolate cell populations showing stable expression of GFP-LC3.

### 2.11. Immunohistochemistry analysis

Immunohistochemical analysis was conducted by staining 3 μm sections, obtained with a RM2125 RTS microtome (Leica Biosystems, Wetzlar, Germany) using anti-LC3-1/2 antibody (diluted 1:300) and anti-ATG-3 (diluted 1:200). The sections were exposed for approximately 12 h at 4°C and then stained with secondary antibodies, along with an avidin-biotin peroxidase complex according to the manufacturer’s protocol (Abcam, Cambridge, UK). An immunoglobulin NC was used to avoid nonspecific binding. The ATG12 and LC3 immunostained the cytoplasm of tumour cells. The staining was regarded as negative when the extent of staining was < 10%. The scoring pattern was as follows: > 10% was regarded as weak staining and was ranked as 1+, ≥ 10% was regarded as moderate staining and was ranked as 2+, and > 10% was ranked as 3+ and regarded as strong staining. Overall, negative immunostaining was defined as a score of 0-1, and positive immunostaining as a score of 2-3.

### 2.12. Tumour studies using nude mice and irradiation

BALB/nude mice were injected subcutaneously with 200 μL of 2 × 10^6^ cells in the right axilla region. After injecting with cancerous cells, the developed tumours were subjected to radiation therapy (^60^Co, 10 Gy/min) after the tumours reached a volume of 500 mm^3^. The dimensions of the tumours (width and length in mm) were measured, along with the volume, every 7 days using callipers. The animals were sacrificed after the tumour diameter reached 15 cm, and the tumours were dissected and weighed.

### 2.13. In situ hybridisation of PC tissues

In situ hybridisation was conducted to determine the expression levels of miR-590-5p in PC tissues. For this purpose, 3 μm tissue sections were obtained and processed by deparaffinisation, followed by rehydration. The sections were then fixed with paraformaldehyde. The sections were reconstituted using hybridisation solution followed by incubation for 20 h at 68°C with DIG-labelled locked nucleic acid (LNA) probes, or with scrambled RNA as an NC. The sections were further incubated along with anti-digoxigenin () alkaline phosphate conjugates, detected using nitroblue tetra-zolium/5-bromo-4-chloro-3-indolyl phosphate (NBT/BCIP); then, the slides were counterstained with Nuclear Fast Red.

### 2.14. Statistical analysis

All results are expressed as the mean ± SD for experiments performed at least in triplicate (n = 3). Statistical evaluation was performed using analysis of variance and Student’s *t*-test. A value of P < 0.05 was regarded as significant.

## 3. Results

### 3.1. The miR-590-5p was differentially expressed in radioresistant PC cell lines

We confirmed and established the radioresistance of the AsPC-1 and Capan-2 human PC cell lines. The cell lines were exposed to programmed irradiation of ^60^Co, as previously described. The cell lines were named accordingly as AsPC-1/RR1, AsPC-1/RR2, Capan-2/RR1, and Capan-2/RR2. A clonogenic survival assay was used to compare the radiosensitivity of radioresistant cell sublines against the parental or non-resistant cancer cell lines, and the fraction of colonies that survived after irradiation were determined. The radioresistant cell sublines showed significantly enhanced resistance to ionising radiation compared to the parental cells (Figure 1A and B). However, the results of the colony formation and cell proliferation assays showed that both the radioresistant cells and the parental cells showed similar proliferations when studied *in vitro* (Figure 1B and C). Table 4 lists the radiosensitivity of the selected radioresistant cell lines. In addition, microarray analysis showed that approximately 21 miRs were differentially expressed in the radioresistant AsPC-1 cell sublines compared to the respective parental cell lines (Table 5). RT-PCR analysis confirmed the results were in agreement with the microarray analysis (Table 5), suggesting that both sublines of radioresistant cells showed decreased expression of miR-590-5p compared to their parental cell lines (Figure 1D).

**Figure 1:**
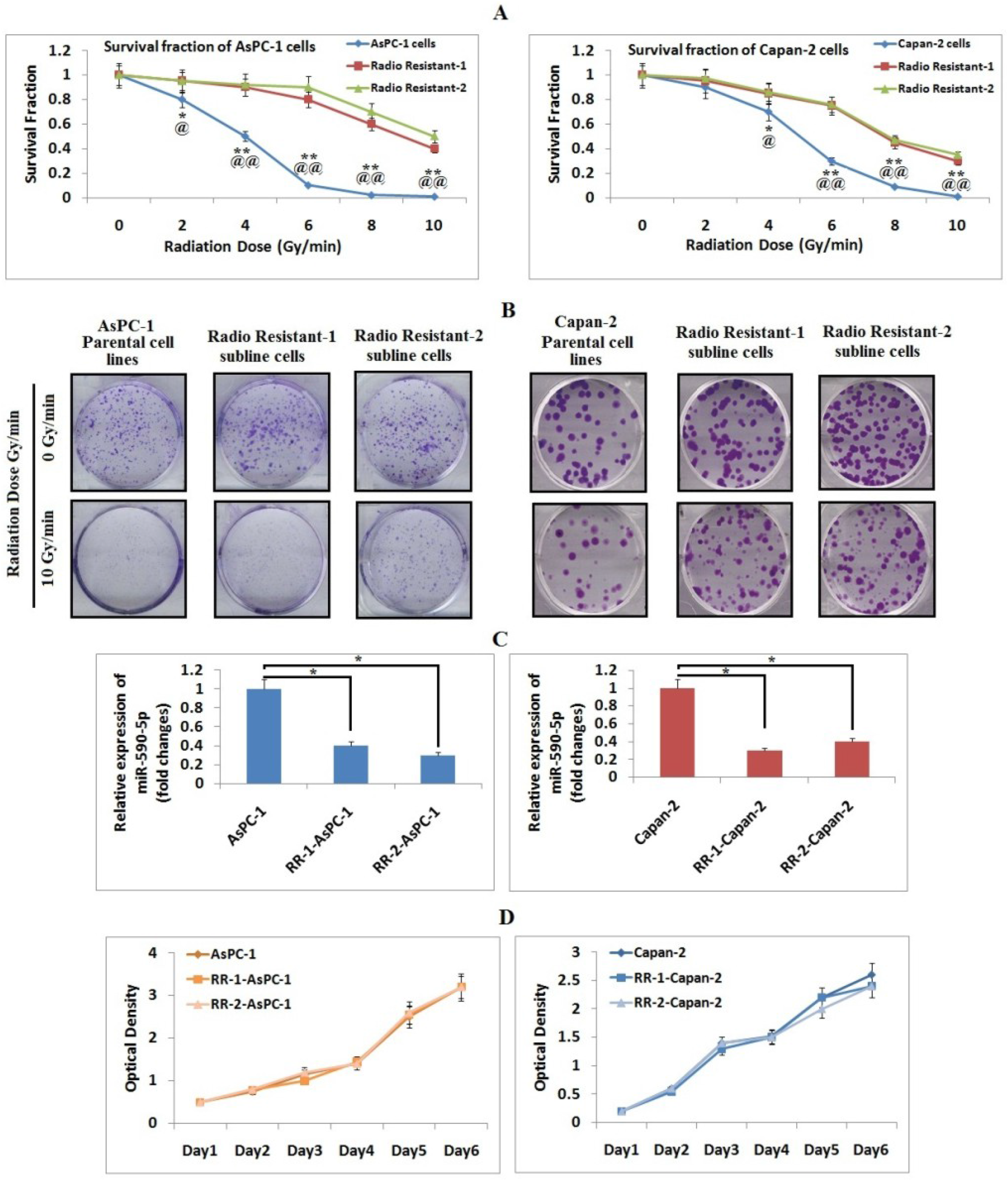
miRNA expression profile of radio resistant pancreatic cancer cells. **A**: Survival fraction of radio resistant PC cells along with their parental cells post exposure to radiation of 0, 2, 4, 6, 8, or 10 Gy. The results are calculated as the means ±SD of values (n=3). *P < 0.05 compared to Radio resistant-1. ^@^*P* compared to Radio resistant-2, **P < 0.01 compared to Radio resistant-1. ^@@^*P* compared to Radio resistant-2. **B**: Crystal violet staining of cell colonies of radio resistant AsPC-1 and Capan-2 along with their respective parental cells, 2 weeks post radiation of 0, 2, 4, 6, 8, or 10 Gy/min. C: Real time PCR (RT-PCR) expression of miR-590-5p in PC cell lines, U6 was used as control. Statistical significance between the groups was done by ANOVA. The results are mean ±SD of values (n=3). *P < 0.05 compared to respective RR cell lines.

**Table 4:**
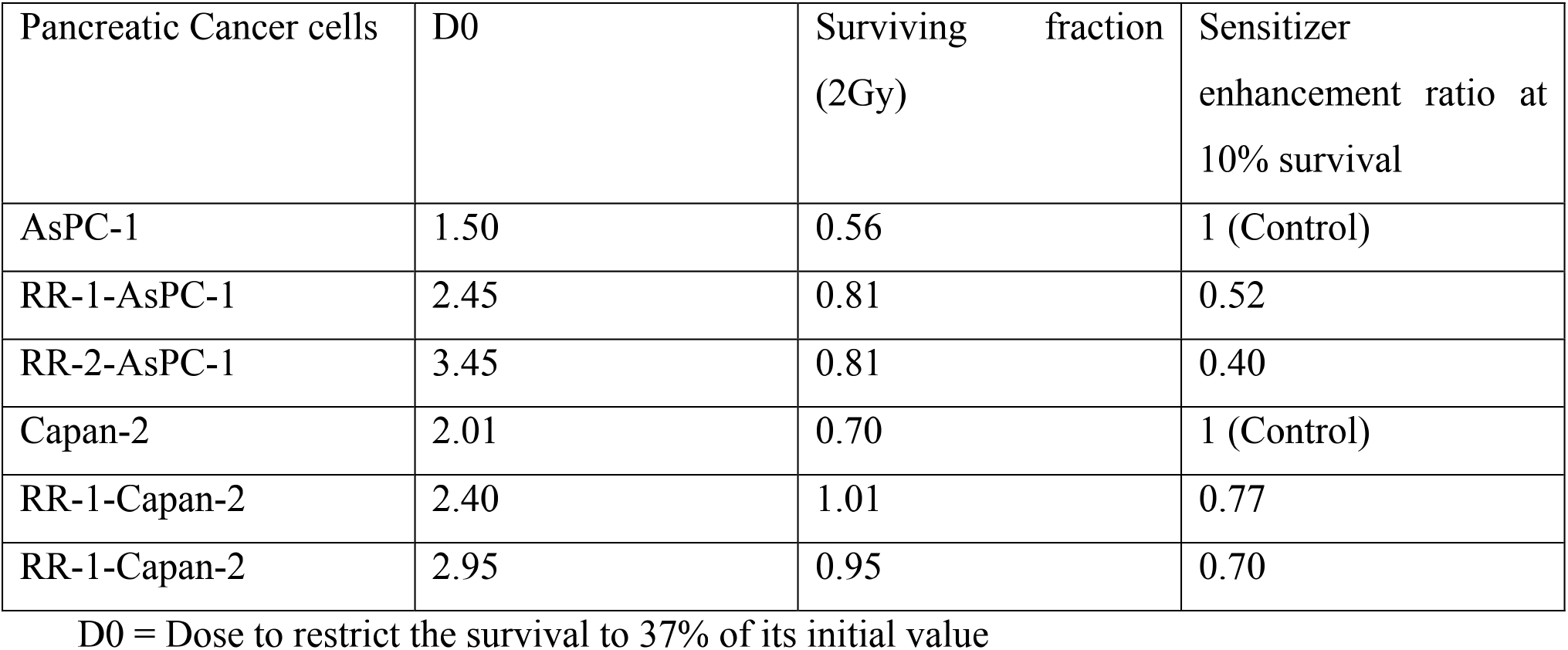
Parameters for radio sensitivity for PC cells and sublines.

**Table 5:**
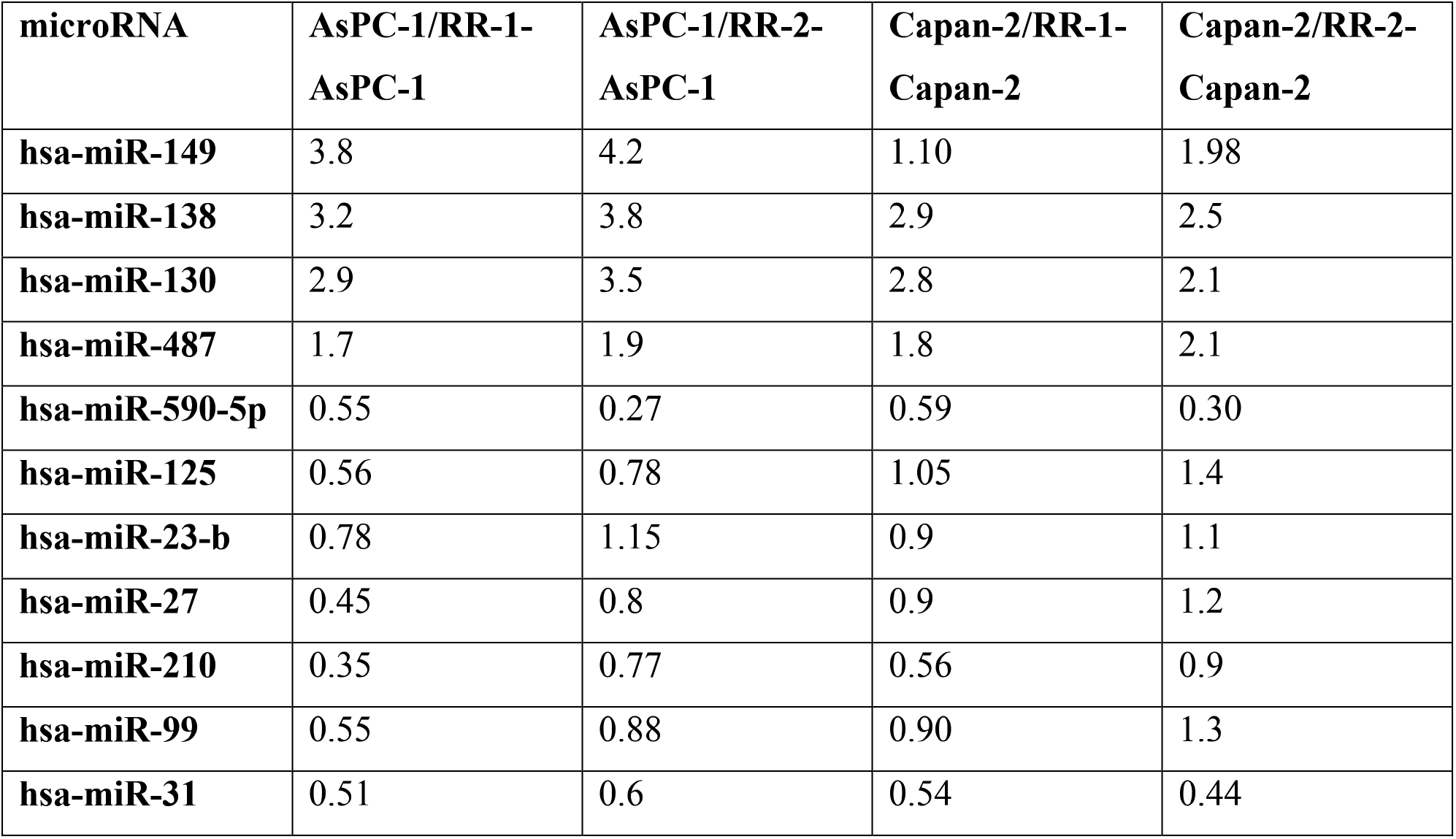
Relative expression of microRNAs in PC cells and sublines, results are fold changes compared to parental cells.

### 3.2. Radioresistant cancer subline cells showed increased autophagy

Autophagy is a protective response occurring in cancer cells during radiation therapy to protect them from radiation-mediated damage, which results in radioresistance [14,15]. We postulated that the radioresistant PC subline cells showed elevated autophagy, and evaluated autophagosomes (in both the radioresistant PC subline cells and their parental cells). Electron microscopy showed that the formation of autophagosomes increased significantly in radioresistant subline cells (Figure 2A). The GFP-LC3-expressing radioresistant PC cells showed a shif in signals, which is indicative of the formation of autophagic vacuoles (Figure 2B), and increased expression of the LC3-2 protein was associated with downregulation of p62 in radioresistant PC cells (Figure 2C). To confirm the involvement of autophagy in radioresistant cell lines, we treated the subline cells with the autophagolysosomal inhibitor chloroquine (CLQ), and observed the effect on levels of LC3-2 proteins. The results showed that CLQ treatment decreased autophagy in radioresistant cells (Figure 2D) and sensitised the PC cells to ionising radiation (Figure 2E). However, treatment with CLQ did not alter the results of the colony formation assay without ionising radiation (Figure 2F). Taken together, the results confirmed that autophagy was involved in the radioresistance of PC cells.

**Figure 2:**
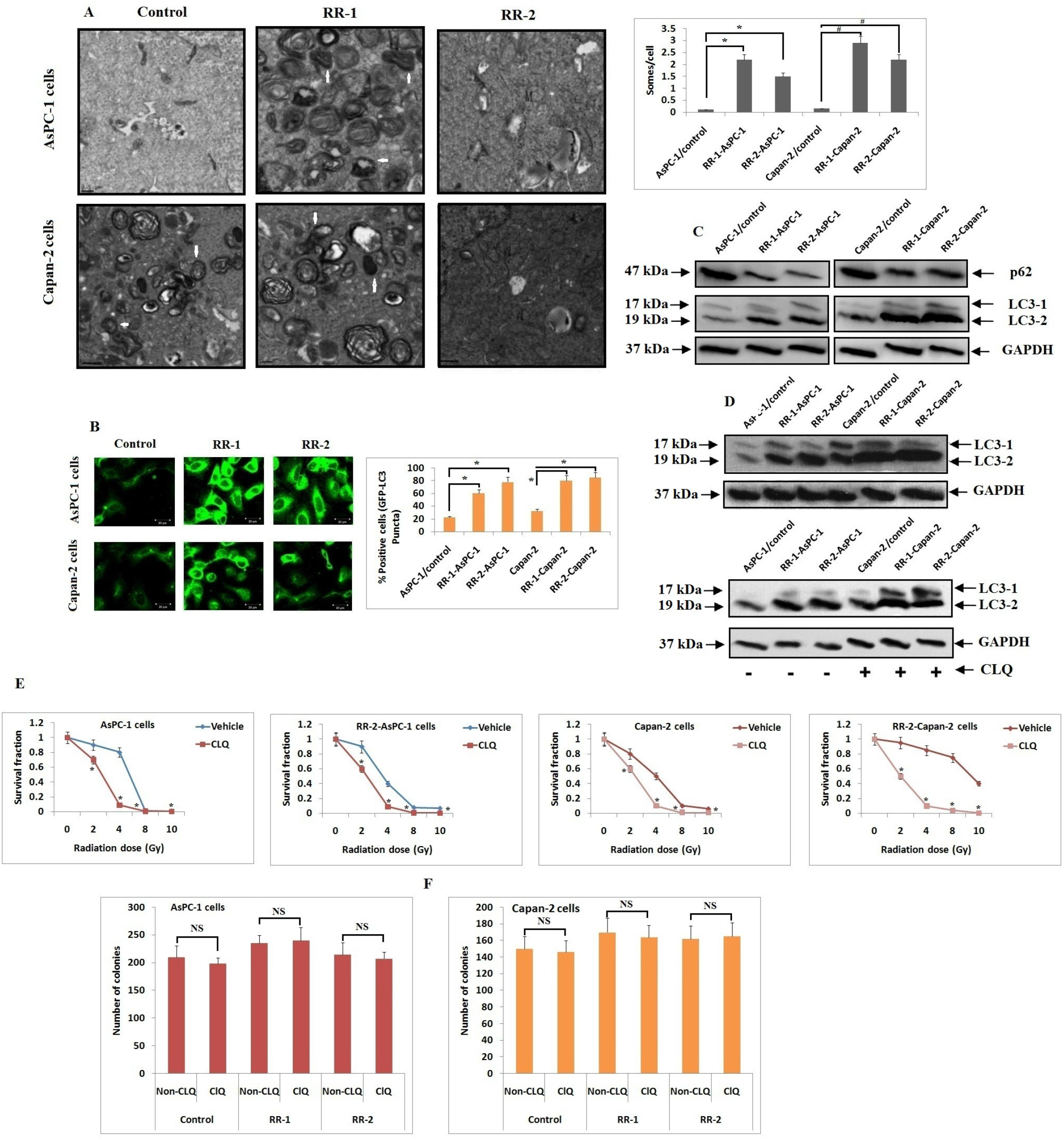
Radio resistant PC cells exhibit elevated autophagy. **A**: Electron microscope was used to assess autophagy in PC cell lines which showed a varying pattern of radiosensitivity (Scale bars, 500 nm). Total numbers of autophagosomes per cross-sectioned cell were counted. The data presented is mean ± SD (n=3). *P < 0.05 compared to control. **B**: The PC cells observed using fluorescent microscope (200X magnification) showed stable expression of GFP-LC3 fusion protein. The extent of autophagy (%) was calculated for positive GFP-LC3 puncta cells. **C**: Immunoblotting study was done to identify formation of autophagosome in cell lysates using LC3 and p62 antibodies. **D**: Radio-resistant PC cells and their parental cell lines were exposed to CLQ (10 μM) for 24 hours prior submitting the samples to immunoblotting study for expression of LC3 **E**: The PC cells were exposed to CLQ for 24 hours prior to radiation, the survival fractions were evaluated. The data presented is mean ± SD (n=3). \**P* < 0.05 compared to vehicle treated cells. **F**: Effect of treatment of CLQ on formation of colonies. The cells were exposed to CLQ for 7 days. The data is presented as mean ± SD (n=3).(NS = Not Significant).

### 3.3. The miR-590-5p blocked autophagy in radioresistant PC cells

We next postulated that deregulation of miRs led to increased autophagy. To test this possibility, the AsPC-1/RR2 and Capan-2/RR2 subline cells were transfected with miR mimics or inhibitors for each of the 13 miRs, in accordance with the expression patterns shown in semi-quantitative PCR and later found during autophagy. Upregulation of either miR-590-5p or miR-18a in radioresistant AsPc-1/RR2 subline cells inhibited the conversion of LC3-1 to LC3-2 (Figure 3A). The results suggested that miR-590-5p blocked the suppression in both radioresistant cell lines compared to their parental cell lines. Because miR-590-5p inhibited the expression levels of both LC3-1 and LC3-2 more so than did miR-18a, we studied the effect of targeted miR-590-5p on autophagy. The radioresistant PC cancer cell lines that showed decreased expression of miR-590-5p were transfected with miR-590-5p mimics to enhance the expression of miR-590-5p. Overexpression of miR-590-5p resulted in a significant increase in the expression of p62, while it suppressed the lipidation of LC3 (Figure 3B). We also found a decrease in the number of autophagosomes in the miR-590-5p-overexpressing cancer cells (Figure 3C and D). Upon treating the cells with CRQ, a significant suppression of autophagic flux was observed in cells showing overexpression of miR-590-5p (Figure 3E). Later studies showed that overexpression of miR-590-5p, which resulted in inhibition of autophagy, resulted in radiation-mediated cytochrome c release and activation of caspase 3 and caspase 9, indicating that miR-590-5p activated a radiation-induced mitochondrial apoptosis cascade (Figure 3F). Overall, the results suggested that miR-590-5p suppressed autophagy in PC cells.

**Figure 3:**
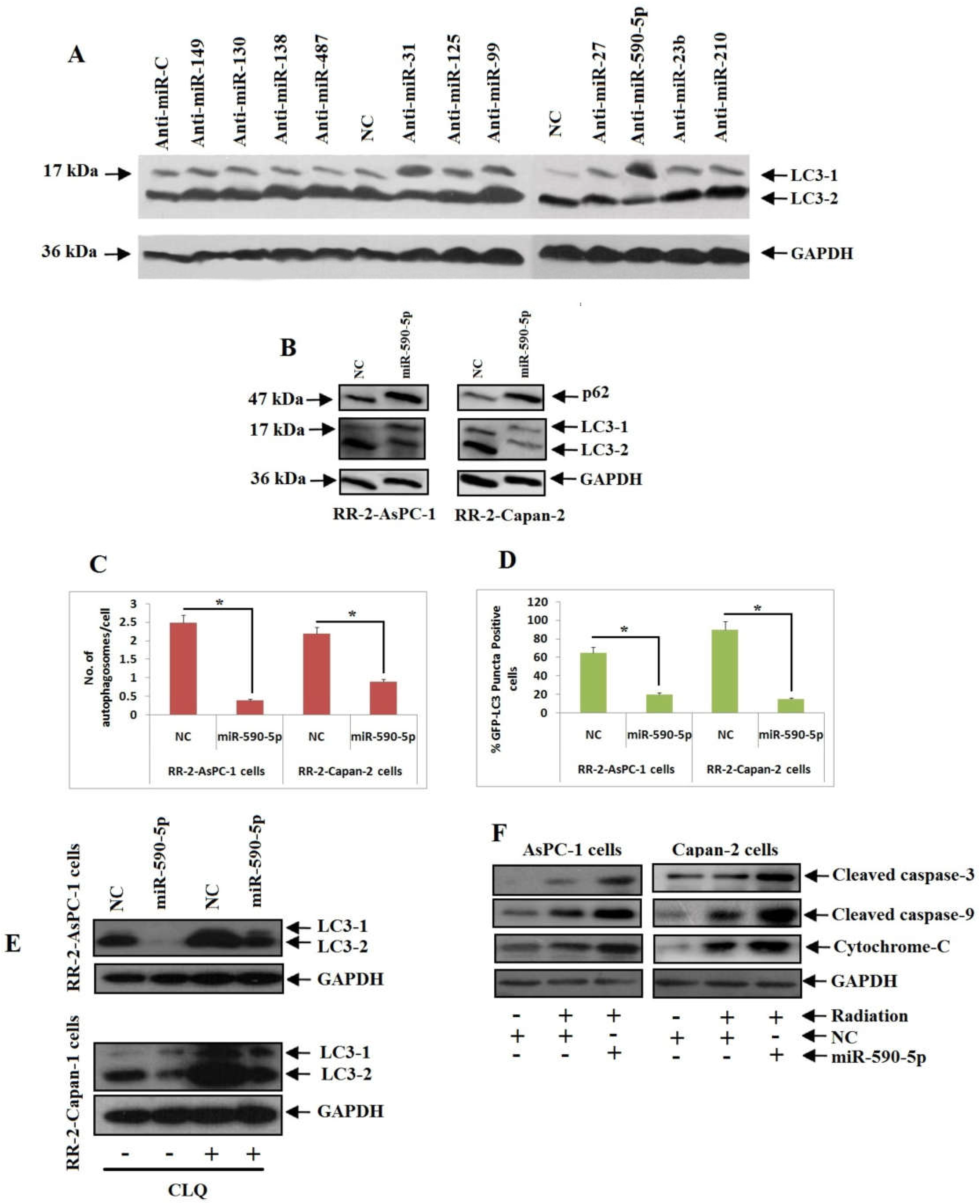
miR-590-5p inhibited autophagy in PC cells. **A**: Radio resistant-2 AsPC-1 cells received transfection of miRNA mimics or inhibitors. The expression of LC3-1 and LC3-2 was identified by Immunoblotting studies. **B**: Radio resistant PC cells received transfection of miR-590-5p mimics or NC and expression of LC3-1 and LC3-2 was studied by Immunoblotting studies after 48 hours of transfection. **C**: The total number of autophagosomes per cell was studied using electron microscopy. *P < 0.05 compared to NC. **D**: The radio-resistant PC cells received transfection of miR-590-5p mimics or NC and after 48 hours were submitted to fluorescent microscopy, \**P* < 0.05 compared to NC. **E**: Immunoblotting analysis for expression of LC3-1 and LC3-2 of radio resistant PC cells transfected with miR-590-5p mimics or NC and after 48 hours followed by exposure to CLQ for 24 hours. **F**: The PC Cells received transfection of miR-590-5p, NC and 48 hours after transfection received radiation of 4 Gy/min. After 24 hours of radiation the proteins were isolated and subjected to Immunoblotting study.

### 3.4. The miR-590-5p enhanced the radiosensitivity of PC cell lines

Based on the decrease in levels of miR-590-5p in radioresistant cell lines, we predicted that miR-590-5p would enhance the sensitivity of PC cancer cells (Figure 4A). The results showed that miR-590-5p mimics significantly enhanced the frequency of radiation-mediated deaths in resistant subline cells (Figure 4B). The radiation parameters have been previously described [16]. The results showed that the value of D0, i.e. the dose required to decrease survival to 37% for AsPC-1/RR2 cells transfected with scrambled mimics, was 1.66, whereas cells transfected with miR-590-5p mimics showed a significant decrease to 1.05. The SER10 values for AsPc-1/RR2 cells transfected with miR-590-5p mimics was 1.85, suggesting that miR-590-5p may have a radiosensitising effect. We also found that the effects of miR-590-5p on radiosensitivity did not alter the proliferation of PC cells. Overall, the results indicated that miR-590-5p enhanced the sensitivity of PC cells towards radiation.

**Figure 4:**
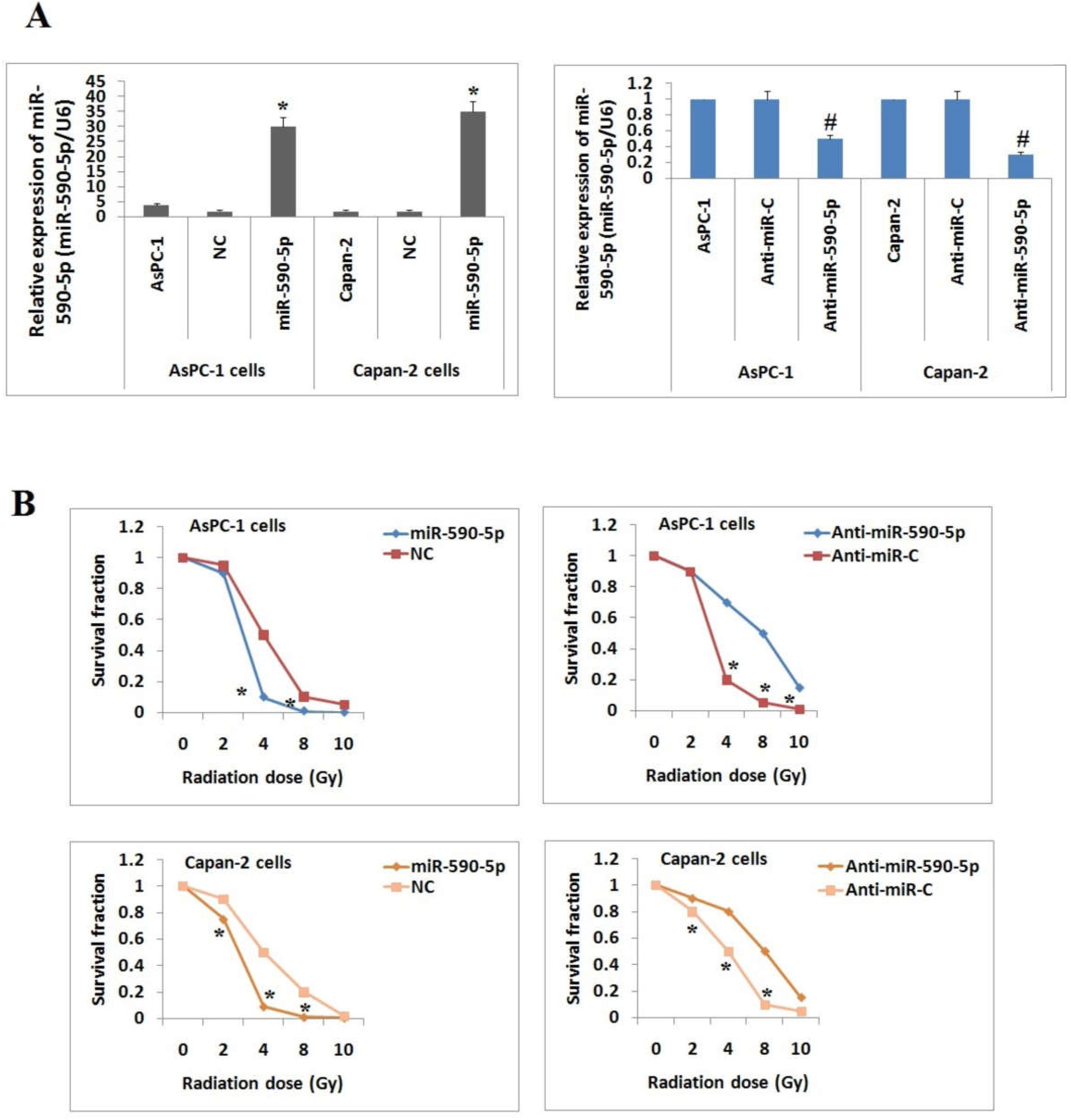
miR-590-5p enhances the sensitivity of PC cells towards radiation. **A**: The effect of miR-590-5p on formation of cell in AsPC-1 and Capan-2 cells. The data presented is mean ±SD (n=3). *P<0.001 compared to NC. **B**: AsPC-1 or Capan-2 PC cell lines were transfected with miR-590-5p inhibitors, mimics or NC, 48 hours after transfection the cells were treated with 0, 2, 4, 6, 8, or 10 Gy of radiation and survival fraction were recorded. The results were mean ± SD (n=3). \**P*<0.01 compared to NC, #P<0.05 compared to Anti-miR-C.

### 3.5. TG-3 was a target of miR-590-5p

We next identified the target gene, which was involved in miR-590-5p-induced autophagy in PC cells. We studied previous RT-PCR data, which suggested possible autophagic genes, and also used bioinformatic tools, such as the TargetScan prediction algorithm, (http://www.targetscan.org/), for identifying a possible target of miR-590-5p. These tools confirmed that ATG-3 was a target of miR-590-5p. The software predicted that miR-590-5p decreased the levels of ATG-3 by binding to two sites of miR-590-5p in the UTR region of ATG-3. The results also suggested that the expression levels of ATG-3 mRNA and its protein were elevated in radioresistant subline cells (Figure 5A). We found that the expression of miR-590-5p was inversely correlated with the expression of ATG-3 mRNA in PC tissue samples, as determined using Spearman’s correlation analysis (Figure 5B). The dual luciferase assay was performed to confirm ATG-3 as the direct target of miR-590-5p; expression of miR-590-5p resulted in a significant suppression of firefly luciferase activity in the 3’-UTR of WT ATG-3, but not in the mutant 3’-UTR of PC cells (Figure 5C). The PC cells were transfected with miR-590-5p mimics, inhibitors, or NC, and the levels of ATG-3 were then estimated using RT-PCR and immunoblotting, which showed that treatment with miR-590-5p resulted in increased expression of ATG-3 (Figure 5D and E). To determine whether miR-590-5p blocked autophagy via suppressing ATG-3, the AsPc-1 cells were transfected with siATG-3 48 h prior to exposing them to radiation. The inhibition of ATG-3 resulted in blockade of radiation-induced autophagy (Figure 5F). Together, the results suggested that ATG-3 was a favourable target of miR-590-5p, and that treatment with miR-590-5p inhibited radiation-mediated autophagy by blocking ATG-3.

**Figure 5:**
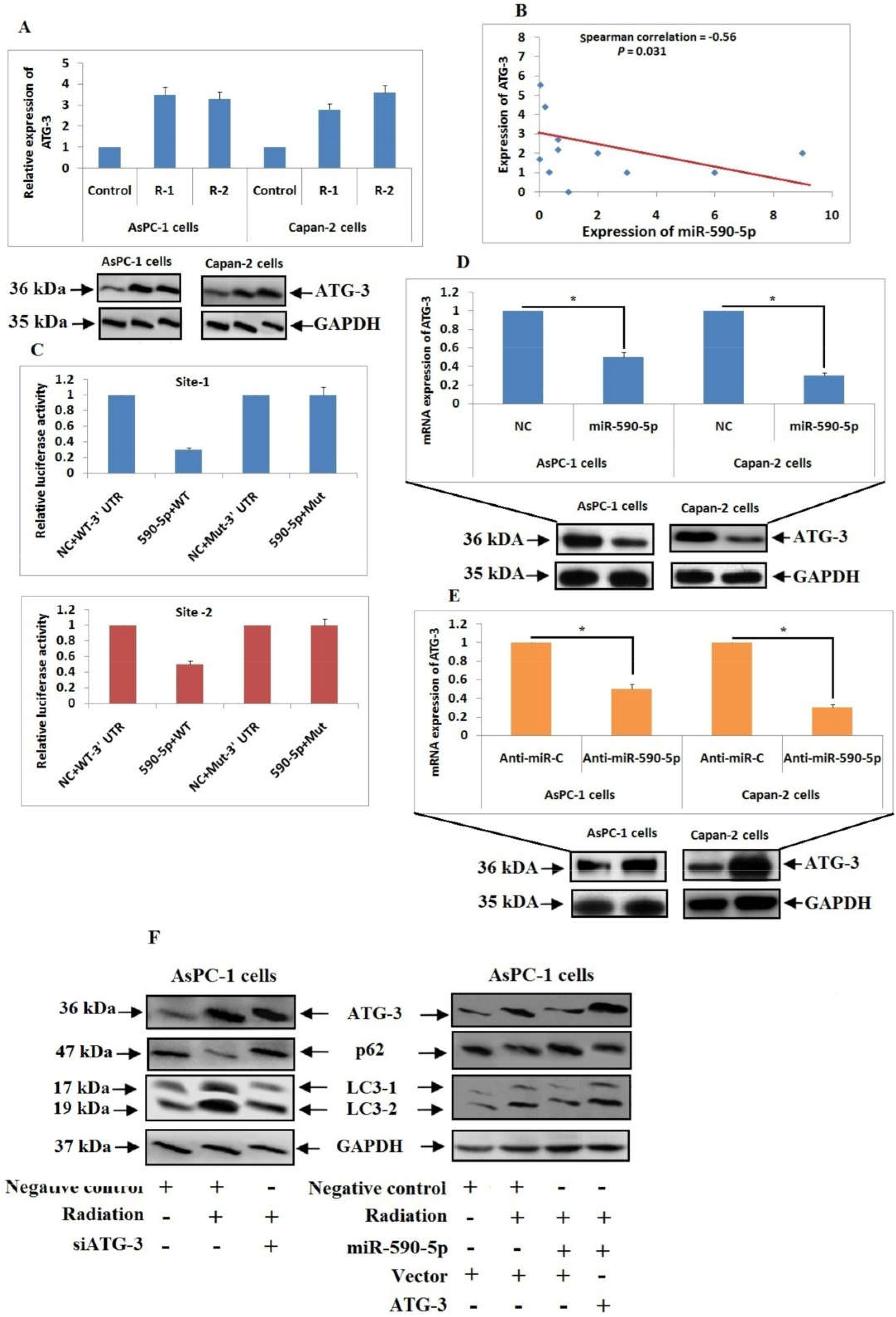
miR-590-5p directly targets ATG-3. **A**: Relative expression of ATG-3 in PC cells using RT-PCR and Immunoblotting analysis. **B**: Spearman correlation analysis was done to establish correlation between miR-590-5p and ATG-3 mRNA expression in tissue samples of patients with PC (n=10). C: Luciferase assay using a luciferase reporter with mutant (mut) or wild-type (WT) ATG-3 3’UTRs were done after co-transfection of miR-590-5p mimics or NC. NC was used for normalization of data and was set as 1 (\**P* < 0.05 compared to NC+WT 3’ UTR). **D and E**: The PC cells received transfection with miR-590-5p mimics or inhibitor or NC. The expression of ATG3 was determined using RT-PCR and Immunoblotting study. The statistical significance was established using ANOVA and Student t tests \**P* <0.05 compared to NC and Anti-miR-C respectively. **F**: AsPC-1 cells were transfected with siATG3, empty vector, NC, miR-590-5p or ATG3, followed by exposure to radiation, 24 hours post radiation after radiation expression of LC3-2, ATG3 and p62 were examined by Immunoblotting study.

### 3.6. The expression of miR-590-5p enhanced the radiosensitivity of PC cells via blocking radiation-mediated autophagy, and was inversely correlated with human PC tissue

Based on the involvement of miR-590-5p in inhibiting radiation-induced autophagy via blocking ATG-3, we extended our study to confirm that miR-590-5p-mediated radiosensitisation was ATG-3 dependent. We found that the inhibition of ATG-3 caused a significant increase in the sensitivity of PC cells to radiation (Figure 6A), and that this inhibition was inhibited by overexpression of ATG-3 (Figure 6A). We also confirmed the role of miR-590-5p and its potential target, ATG-3, in *in vivo* sensitivity using tumour xenografts in mice. The results of the *in vivo* study suggested that upregulation of miR-590-5p resulted in a significant increase in the radiosensitivity of induced tumours of AsPc-1 cells upon exposure to radiation in mice (Figure 6B). We also found that blocking the expression of ATG-3 enhanced the sensitivity of AsPc-1-induced tumours towards radiation treatment in mice (Figure 6C). When we analysed tumours 24 h after a single dose of 10 Gy/min, the expression of miR-590-5p resulted in inhibition of radiation-induced autophagy in a similar manner as seen in the *in vitro* experiments (Figure 6D). The results suggested that upregulation of miR-590-5p in mice resulted in an increase in radiation-induced release of cytochrome C and upregulation of caspase 3 and caspase 9 (Figure 6E). CLQ did not synergise with the inhibition of autophagy mediated by miR-590-5p after exposure to radiation, but CLQ caused a significant elevation in radiosensitivity upon suppression of miR-590-5p using an anti-mir-590-5p inhibitor (Figure 6F).

**Figure 6:**
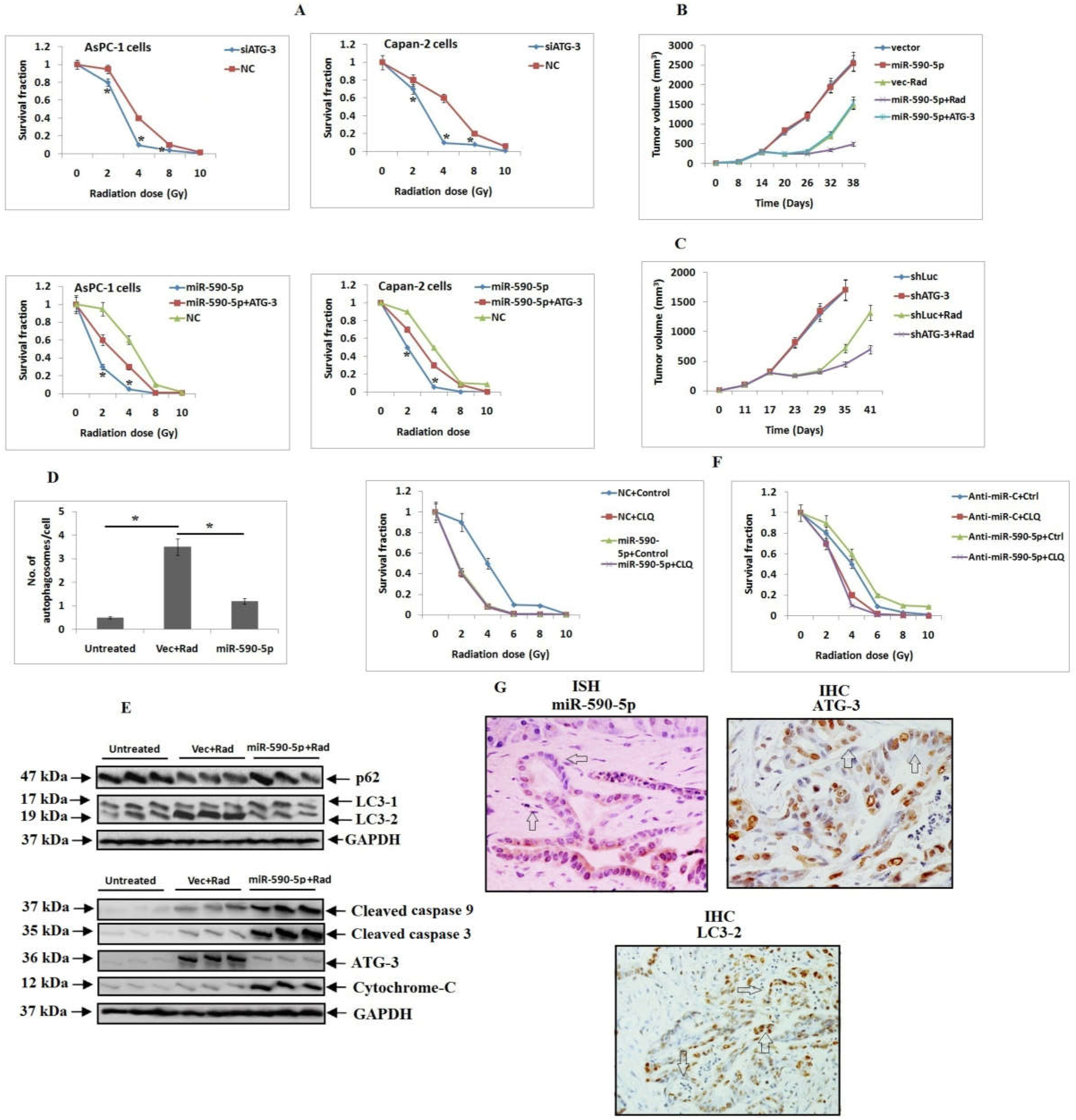
miR-590-5p enhances the sensitivity of PC cells to radiation via inhibiting radiation-mediated autophagy. **A**: The PC cell lines received transfection of siATG3, miR-590-5p, NC, or ATG3 followed by exposure to radiation of predefined Gy. The results of survival fraction were estimated for each group as the mean ±SD (n=3). Statistical significance was established using ANOVA and Student t tests *P* <0.05 compared to NC (*) and miR-590-5p+ATG-3(#). **B**: Tumor volumes of nude mice injected subcutaneously with AsPC-1 cells transfected with miR-590-5p over-expressing vector, lentiviral vector or an ATG-3 expressing vector lacking 3’UTR. The tumors were exposed to radiation (single dose 10 Gy/min) when the volume reached 500 mm^3^. **C**: The nude mice were injected subcutaneously with AsPC-1 cells transfected with control lentivector (ShLuc) or shATG-3 vector transfected cells, the tumor volumes were recorded. **D**: Electron microscopy was used to evaluate autophagy in cells isolated from tumor xenografts samples, *P<0.05. **E**: Proteins were isolated from cells of tumor xenografts samples and proteins were subjected to Immunoblotting study. F: The PC cells were transfected with miR-590-5p, anti-miR-590-5p inhibitor or NC, the cells were maintained in presence or absence of CLQ prior to subjecting them to radiation. **G**: The mature miR-590-5p in human pancreatic cancer tissue specimen was identified opting In situ hybridization (ISH) (n=54). Immunohistochemistry (IHC) was done using antibodies for ATG-3, LC3.

We also established a correlation between the expression of miR-590-5p and autophagy in PC tissues, by using a LNA-miRNA probe and in situ hybridisation to determine the levels of mature miR-590-5p in cancer tissue samples. In addition, immunohistochemical analyses of tissue sections of human PC using ATG-3- and LC3-1/2-specific antibodies suggested that the clinicopathological factors of pancreatic tumours showed a weak correlation (low expression) with ATG-3 and LC3 (Table 1), with increased expression of miR-590-5p (Figure 6G) (Table 6). Together, these results confirmed that miR-590-5p enhanced the sensitivity of PC cells to radiation via inhibiting radiation-induced autophagy, and further confirmed that the expression of miR-590-5p was inversely associated with human PC tissue.

**Table 6:**
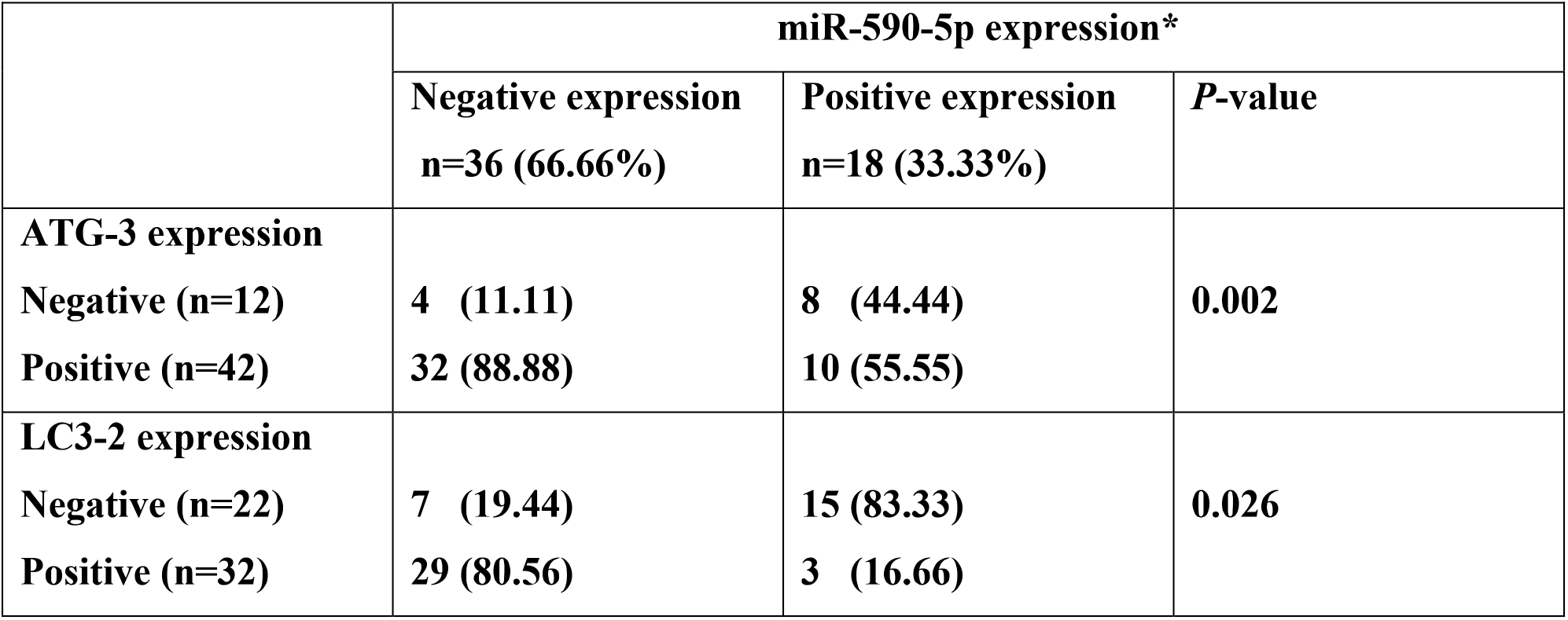
Data showing correlation between expression of miR-590-5p, ATG-3 and LC3-2 in pancreatic cancer tissue specimens, the expression of miR-590-5p was identified by in situ hybridization. Expression of ATG-3 and LC3-2 was done by immunohistochemical analysis. *P* values were calculated by Pearson x^2^ test.

## 4. Discussion

In the present study, we identified miR-590-5p as a blocker of autophagy. We confirmed that ATG-3 was a target of miR-590-5p, which was also affected by radiation-mediated autophagy. We also confirmed that miR-590-5p enhanced the sensitivity of PC cells against radiation by inhibiting autophagy. The novel results of our study strongly suggested miR-590-5p played a major role in regulating autophagy, and confirmed that miR-590-5p enhanced the radiosensitivity of PC cells against radiation via inhibiting radiation-mediated autophagy.

Autophagy has been reported to be crucial for maintaining homeostasis at the cellular level; it plays a protective role in cells during periods of low nutrition and has also been reported to have a dual role in cancer [14–16]. The absence of autophagic genes can lead to increased tumorigenesis [17], and previous studies have reported that autophagy contributes to tumour progression during chemotherapy [18–20]. Previous studies have also shown that modalities for treating cancer may promote autophagy, both *in vitro* and *in vivo* [21,22]. Radiation-mediated autophagy defends cells against radiation-induced damage, to enhance radioresistance [21,18]. The results of the present study confirmed that autophagy is a form of resistance against radiotherapy, and suggested approaches to target autophagy, which is involved in the radioresistance to cancer treatment.

There has recently been increased interest in approaches involving mRNA expression in tumours, as well as studies of sensitivity to chemotherapy or radiation therapy as a method of modulating cellular sensitivity [23]. Resistance is always challenging when treating cancer, and it is one of the major challenges in treating PC [24]; thus, identifying the mechanisms and pathways involved in radiosensitivity is crucially important in the control of this type of cancer. In the present study, we identified approximately 13 differentially expressed miRs that correlated with radioresistance.

Among them, miR-590-5p was downregulated to a large extent in radioresistant cell lines compared to their parental cells. Previous reports have suggested that miR-590-5p is differentially regulated in hepatic, cervical, gastric, breast, and renal cancers [25–28], although the exact role of miR-590-5p remains unclear. In addition, a previous study confirmed its role in enhancing chemosensitivity [29] and radiosensitivity [30]. However, the exact role of miR-590-5p in PC, and its associated chemoresistance, remains unexplored and needs to be characterised.

In the present report, we studied the effect of miR-590-5p on PC cells, and the effect of miR-590-5p on the radiosensitivity of these cells, showing that miR-590-5p significantly enhanced the sensitivity of PC cells. It has been confirmed that cancer cells need autophagy for survival under stressful conditions, such as exposure to chemotherapy and radiotherapy, nutrient deprivation, hypoxia, and metabolic stress [31–34]. Inhibition of autophagy enhances the potential of treatment independent of proliferation [31], because PC cells possess increased autophagy compared to all other cancer cell lines [35]. Recent reports have shown that autophagy is vital for the growth of PC cells [36–37], and it is also a mechanism protecting starving cells. Based on these findings, inhibiting autophagy could be a possible cancer treatment. However, it is still unclear whether only inhibiting cell proliferation or the fact that autophagy facilitates the survival of PC cells under the situations which occur during the tumour growth and proliferation. Therefore, the pathways and mechanisms regulating autophagy need to be identified. In the present study, we identified autophagy as a possible target for treating PC. Our findings confirmed the role of autophagy in radioresistance, and further showed that miR-590-5p inhibited, and even enhanced, the sensitivity of PC cells to radiotherapy. Our findings also showed that exposing radioresistant PC cells to CLQ (an autophagy inhibitor) did not synergistically contribute to inhibiting the autophagy induced by miR-590-5p after radiation therapy. However, by suppression of miR-590-5p using an inhibitor, we found that CLQ significantly enhanced the sensitivity; our results therefore suggested that expression of miR-590-5p was associated with the radiation sensitivity of PC cells. Thus, an advance analysis of patients with respect to levels of miR-590-5p prior to submitting them for treatment could be advantageous and may improve the overall prognosis.

## Conclusion

The present study showed that radioresistant PC cells exhibited increased autophagy, and treatment with miR-590-5p resulted in suppression of autophagy via targeting ATG-3, thereby decreasing autophagy and improving radiosensitivity. Our findings confirmed the role of miR-590-5p in regulating autophagy, and suggested a new pathway that may significantly improve therapies for PC.

## Declaration

The authors have no conflict of interest.

